# Greening rates are sensitive to methodology and biology; comment to Sustained greening of the Antarctic Peninsula observed from satellites

**DOI:** 10.1101/2024.11.07.622227

**Authors:** S. Bokhorst, S. Huisman, I.K. de Jonge, T.A.J. Janssen, J.H.C. Cornelissen, K. A. Hughes, P. Convey

## Abstract

Roland et al.^1^ claim to provide evidence for a ‘greening trend’ throughout the Antarctic Peninsula region over the last four decades, based on satellite remote sensing data. However, the early period vegetation cover estimates do not match with the likely extent of vegetated areas in this region at that time, raising doubts about the magnitude of any greening trend. Furthermore, growth rates of mosses and higher plants in Antarctica are insufficient to explain the 14-fold green cover expansion claimed, and neither have such changes been observed at long-term monitoring sites or from field warming studies. The reported satellite time series analyses of the presented trend seems biased by satellite image availability, lack of consistency in the areas covered by imagery and processing pitfalls. Antarctic terrestrial ecosystems are indeed predicted to become greener with climate change, but at much slower rates than reported by Roland et al.^1^.

## Field observations

Ground-truthing of satellite imagery is crucial to validate remotely sensed data, as also argued by Roland et al.^1^. While robust ground observations of Antarctic vegetation are scarce, they do exist, and could have been used to align satellite imagery to known distribution and extent of moss/higher plant cover. The reported green cover (NDVI > 0.2) for the studied region for 1986 was 0.863 km^2^, but this is very low compared to botanical reports from the 1960s to 1990s. For instance, vegetation cover on several islands in the South Orkney Islands and South Shetland Islands is typically described as extensive, including continuous cover across several hundred meters to kilometers in coastal areas^2,3^. These reports raise the fundamental question of whether the 1980s green cover estimates used by Roland et al.^1^ provide a true reflection of the extent of vegetated areas along the Antarctic Peninsula and Scotia Arc at that time. The 1986 satellite-derived vegetation cover is further questionable as there appears to be no change in green cover between 1986-2021 for the region containing the South Shetland Islands (their Fig. 3e), which encompass Byers Peninsula (Livingston Island) where current green vegetation cover reaches 8 km^2^ (ASPA 126), which is 10 times the satellite-derived estimate for the entire Antarctic Peninsula, and highlights the complexity of matching satellite data with vegetation ground cover.

True moss banks are illustrated as a potential source of the reported expansion by Roland et al.^1^ (their Fig. 1). These banks have existed along the Antarctic Peninsula for thousands of years^2^, and show enhanced vertical growth under warmer and wetter conditions^4^, but there are no reports of extensive horizontal expansion, and many such banks have distinct vertical edges. Studies on recently deglaciated terrain (since the 1950s) report very low values of vegetation cover (0.4 hectares) across sites on the South Shetland Islands, one of the greenest and floristically richest parts of the Maritime Antarctic^5,6^. If a 14-fold moss cover expansion had driven the observed greening trend, it would have been recorded at such sites. However, no such scale of expansion has been documented, including on islands in Ryder Bay (Adelaide Island, c. 68ºS) where our research group and collaborators have monitored vegetation since 2004 (personal observations SB; Bokhorst et al.^7^), or on Signy Island (South Orkney Islands), where detailed island-wide ground vegetation mapping of higher plants and bank-forming mosses has taken place across time (1960s, 2000s and late 2010s)^8^. Enhanced sporulation, which could support moss expansion, occurs under field warming^9^, but moss expansion was not recorded in field warming studies on Signy Island between 2003 and 2013^10^. Rather, rapid microalgal colonization of soils in field warming studies on Signy Island (60° S) and Alexander Island (70-71° S) has been documented^11^, suggesting that algae can respond rapidly to climate warming. We suggest this process could apply across much of the Antarctic Peninsula region, where mats of the green alga *Prasiola crispa* can cover hundreds of square meters^12^, particularly under the influence of penguin colonies and seal aggregations. While the spatial extent of these mats can change widely between years and even days, resulting from water availability and wind dispersal, even the relatively rapid growth rates of algae alone are unlikely to account for the large inter-annual variability in green cover reported by Roland et al.^1^.

**Figure 1.**
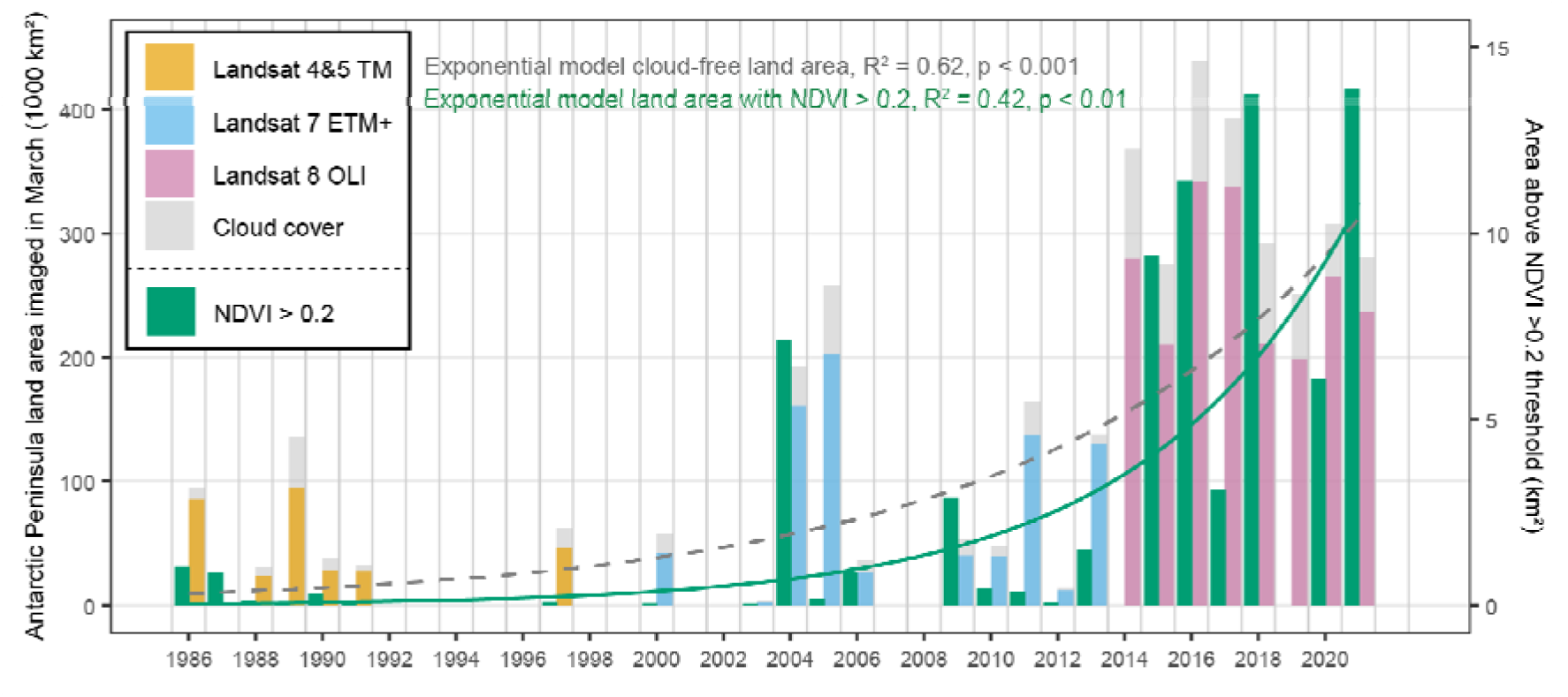
Changes in the extent of imaged land area and land area with NDVI above the 0.2 threshold. The data only include images from the month of March, with the highest quality assurance and with less than 40% cloud cover over the Antarctic Peninsula between 1986 and 2021 (sensu Roland et al.^1^). The dark green bars represent the land surface area with NDVI above the 0.2 threshold and was retrieved from Figure 2 in Roland et al^1^. The land area obstructed by clouds in the coverage of the three Landsat sensors are indicated in grey.

## Ecophysiology and growth

Although Antarctic greening is predicted, and expansion by the two native vascular plants and mosses have been carefully documented at sites along the Antarctic Peninsula and Scotia Arc^8,13^, the scale of these expansions remains far too small to explain the large-scale patterns reported^1^. Attributing the increase solely to moss expansion is not plausible from an ecophysiological perspective. Antarctic mosses grow slowly, typically at 1-5 mm/year, and mostly by vertical shoot growth^4^. It is therefore highly unrealistic that moss horizontal expansion can contribute significantly to the scale of the observed increases in green cover over 35 years, particularly the large inter-annual increases and decreases of several square kilometers (of similar magnitude to the total moss area inferred in many years in the study period). Likewise, any abrupt green cover declines would most likely involve complete destruction of the entire moss community which, in turn, cannot recover within one growing season. Although destruction of moss carpets has been linked to increased fur seal numbers, such trampled and over-manured moss carpets are rapidly overgrown by terrestrial algae^14^. If the remote sensing data reflect vegetation cover change, which is uncertain without ground-truthing, we suggest that a more likely biological candidate for the rapid interannual variation may be sought among freshwater and terrestrial algae and microbial mats. Snow algae may similarly contribute, as a large proportion of the up to several square kilometers that can develop in a typical season^15^ is likely to form within the 300 m ‘buffer zone’ incorporated by Roland et al.^1^ in their definition of the boundary of snow and ice-free ground (their supplementary Fig. 2.1). These different algae also contribute to the NDVI signal^15^, and can vary widely on both intra- and interannual timescales as they are not part of the permanent vegetation.

Roland et al.^1^ speculated that the interannual variation in their data may indicate changes in moss health, as this affects reflectance^16^. However, for this to affect green cover across the entire Antarctic Peninsula and South Shetland Islands, a large proportion of the many dozens of moss populations/species, inhabiting various habitats, must be simultaneously negatively affected across more than 1000 km of coastline, followed by complete recovery the next season. Given the diverse habitat requirements of Antarctic mosses^17^, such a consistent response to a single disturbance seems highly unlikely.

## Satellite-image trends

The main concern we have with the remote sensing analysis of Roland et al.^1^ is that their greening trend is expressed as land area above a greenness threshold (in km^2^, see their Fig. 2), while the availability of satellite images, and therefore land area imaged (also in km^2^), varied considerably over the study period. Roland et al.^1^ state that there was a poor correlation between observed land percentage and total area above the greenness threshold (see their supplementary Fig. 3.2), but their data indicate a significant correlation explaining 35% of the variation in land area above the 0.2 NDVI threshold (R^2^ = 0.351, p = 0.0029). This indicates that, during some years with larger available land surface area for analysis, they logically observed a larger area above the greenness thresholds, which raises the question of whether the reported greening trend reflects an actual increase in green cover or that more green areas of the Antarctic Peninsula region are included in the analysis each year. In addition, it is unclear whether the same areas are being compared between years (different parts of the Antarctic Peninsula are compared between years). By using the same selection criteria for obtaining Landsat images as Roland et al.^1^, see method, we find an increasing trend in the extent of imaged cloud-free land area since the 1980s, which is very similar to the increasing trend of land area above the 0.2 NDVI threshold (Fig. 1). Especially since the launch of the Landsat 8 satellite in 2013, the coverage of cloud-free land area in March over the AP increased from only 3.6 Mha per year between 1986-2012 to 37.2 Mha per year between 2013 and 2021, dramatically increasing the likelihood of capturing land area above the greenness threshold. In addition, Roland et al.^1^ noted that there was not a single cloud-free area where the images from Landsat-7 and -8 data overlapped, preventing a regional cross-calibration of these sensors while the transition between these sensors reflected the largest increase in greenness (Fig. 1).

## Concluding remarks

Climate change is altering conditions for life in Antarctica, and additional growth and population expansion of primary producers are expected at sites with sufficient water availability. However, the green cover changes proposed by Roland et al.^1^ do not match with either known historical descriptions of distributions or the known ecophysiology and growth characteristics of Antarctic mosses and vascular plants. We suggest that any Antarctic greening trend, while predicted, is presently unproven and is likely to be much more moderate than claimed. This raises the key questions of whether the green cover estimates for 1986-2012 reflect a considerable underestimation and if the reported trend is not biased by the much higher availability of satellite images during 2013-2021.

## Methods

To identify if the observed greening patterns by Roland et al. (see their Fig. 2) could be associated with extent of cloud-free land area imaged we obtained the metadata of the entire Landsat archive covering the Antarctic Peninsula and South Shetland Islands from 1986-2021. Landsat metadata was retrieved from the Earth Explorer website of the United States Geological Survey (https://earthexplorer.usgs.gov/). We selected images using the same selection criteria as used by Roland et al., using only images from the month of March, with the highest quality assurance number (9) and with less than 40% cloud cover. We calculated total land area imaged (in km^2^) using the Landsat scene corner coordinates provided in the image metadata to create a spatial bounding box in combination with the MODIS land-water mask from 2021 (MOD44W v061) retrieved from NASA’s Land Processes Distributed Active Archive Center (LP DAAC). To estimate the land area imaged that was obstructed by cloud cover, we multiplied the land cloud cover value (%) provided in the metadata with the total land area imaged. All land area imaged was summed for each year, including areas that overlapped spatially because multiple overpasses increase the probability of capturing cloud-free land area as well as capturing a higher NDVI in March. To identify any patterns over time we plotted a trend line through the available cloud-free land area imaged (in km^2^) and the land area with NDVI > 0.2 (also in km^2^) derived from Fig. 2 in Roland et al.

## References

1. Roland, T.P., et al. Sustained greening of the Antarctic Peninsula observed from satellites. Nature Geoscience, (2024).

2. Lindsay, D.C. Vegetation of the South Shetland Islands. Brit. Antarct. Surv. Bull. 25, 59–83 (1971).

3. Allison, J.S. and R.I.L. Smith. The vegetation of Elephant Island, South Shetland Islands. Brit. Antarct. Surv. Bull. 33 & 34, 185–212 (1973).

4. Royles, J., et al. Plants and soil microbes respond to recent warming on the Antarctic Peninsula. Curr. Biol. 23, 1702–1706 (2013).

5. Ruiz-Fernandez, J., M. Oliva, and C. Garcia-Hernandez. Topographic and geomorphologic controls on the distribution of vegetation formations in Elephant Point (Livingston Island, Maritime Antarctica). Science of the Total Environment. 587, 340–349 (2017).

6. Smith, R.I.L. Plant succession and re-exposed moss banks on a deglaciated headland in Arthur Harbour, Anvers Island. Brit. Antarct. Surv. Bull. 51, 193–199 (1982).

7. Bokhorst, S., et al. The effect of environmental change on vascular plant and cryptogam communities from the Falkland Islands and the Maritime Antarctic. BMC Ecol. 7, 15 (2007).

8. Cannone, N., et al. Acceleration of climate warming and plant dynamics in Antarctica. Curr. Biol. 32, 1599-1606.e2 (2022).

9. Casanova-Katny, A., G.A. Torres-Mellado, and S.M. Eppley. Reproductive output of mosses under experimental warming on Fildes Peninsula, King George Island, maritime Antarctica. Revista Chilena de Historia Natural. 89, 13 (2016).

10. Bokhorst, S., et al. Usnea antarctica, an important Antarctic lichen, is vulnerable to aspects of regional environmental change. Polar Biology. 39, 511–521 (2016).

11. Wynn-Williams, D.D. Response of pioneer soil microalgal colonists to environmental change in Antarctica. Microbial Ecology. 31, 177–188 (1996).

12. Longton, R.E. Vegetation in the maritime Antarctic. Philosophical Transactions of the Royal Society of London Series B-Biological Sciences. 252, 213–235 (1967).

13. Fowbert, J.A. and R.I.L. Smith. Rapid population increases in native vascular plants in the Argentine Islands, Antarctic Peninsula. Arct. Alp. Res. 26, 290–296 (1994).

14. Smith, R.I.L. Destruction of Antarctic terrestrial ecosystems by a rapidly increasing fur-seal population. Biological Conservation. 45, 55–72 (1988).

15. Gray, A., et al. Remote sensing reveals Antarctic green snow algae as important terrestrial carbon sink. Nat. Commun. 11, 2527 (2020).

16. Turner, D., et al. Mapping water content in drying Antarctic moss communities using UAS-borne SWIR imaging spectroscopy. Remote Sensing in Ecology and Conservation. 68, 168–179 (2018).

17. Gimingham, C.H. and R.I.L. Smith. Growth form and water relations of mosses in the Maritime Antarctic. Brit. Antarct. Surv. Bull. 25, 1–21 (1971).

